# Algorithmic Learning for Auto-deconvolution of GC-MS Data to Enable Molecular Networking within GNPS

**DOI:** 10.1101/2020.01.13.905091

**Authors:** Alexander A. Aksenov, Ivan Laponogov, Zheng Zhang, Sophie LF Doran, Ilaria Belluomo, Dennis Veselkov, Wout Bittremieux, Louis Felix Nothias, Mélissa Nothias-Esposito, Katherine N. Maloney, Biswapriya B. Misra, Alexey V. Melnik, Kenneth L. Jones, Kathleen Dorrestein, Morgan Panitchpakdi, Madeleine Ernst, Justin J.J. van der Hooft, Mabel Gonzalez, Chiara Carazzone, Adolfo Amézquita, Chris Callewaert, James Morton, Robert Quinn, Amina Bouslimani, Andrea Albarracín Orio, Daniel Petras, Andrea M. Smania, Sneha P. Couvillion, Meagan C. Burnet, Carrie D. Nicora, Erika Zink, Thomas O. Metz, Viatcheslav Artaev, Elizabeth Humston-Fulmer, Rachel Gregor, Michael M. Meijler, Itzhak Mizrahi, Stav Eyal, Brooke Anderson, Rachel Dutton, Raphaël Lugan, Pauline Le Boulch, Yann Guitton, Stephanie Prevost, Audrey Poirier, Gaud Dervilly, Bruno Le Bizec, Aaron Fait, Noga Sikron Persi, Chao Song, Kelem Gashu, Roxana Coras, Monica Guma, Julia Manasson, Jose U. Scher, Dinesh Barupal, Saleh Alseekh, Alisdair Fernie, Reza Mirnezami, Vasilis Vasiliou, Robin Schmid, Roman S. Borisov, Larisa N. Kulikova, Rob Knight, Mingxun Wang, George B Hanna, Pieter C. Dorrestein, Kirill Veselkov

## Abstract

Gas chromatography-mass spectrometry (GC-MS) represents an analytical technique with significant practical societal impact. Spectral deconvolution is an essential step for interpreting GC-MS data. No public GC-MS repositories that also enable repository-scale analysis exist, in part because deconvolution requires significant user input. We therefore engineered a scalable machine learning workflow for the Global Natural Product Social Molecular Networking (GNPS) analysis platform to enable the mass spectrometry community to store, process, share, annotate, compare, and perform molecular networking of GC-MS data. The workflow performs auto-deconvolution of compound fragmentation patterns *via* unsupervised non-negative matrix factorization, using a Fast Fourier Transform-based strategy to overcome scalability limitations. We introduce a “balance score” that quantifies the reproducibility of fragmentation patterns across all samples. We demonstrate the utility of the platform with breathomics analysis applied to the early detection of oesophago-gastric cancer, and by creating the first molecular spatial map of the human volatilome.

## Introduction

Electron ionization gas chromatography-mass spectrometry (GC-MS) is widely used in numerous analytical applications with profound societal impact, including screening for inborn errors of metabolism, toxicological profiling in humans and animals, basic science investigations into biochemical pathways and metabolic flux, understanding of chemoattraction, doping investigations, forensics, food science, chemical ecology, ocean and air quality monitoring, and many routine laboratory tests including cholesterol^1^, vitamin D^2^ and lipid levels^3^. GC-MS is widely adopted because of its key advantages, including low operational cost, excellent chromatographic resolution, reproducibility and ease of use.

In GC-MS, the predominant ionization technique is electron ionization (EI), in which all compounds that elute from the chromatography column are ionized by high energy (70eV) electrons in a highly reproducible fashion to yield a combination of fragment ions. Because fragmentation occurs simultaneously with ionization, an essential computational step in the analysis of all GC-MS data is the “spectral deconvolution” - the process of separating fragmentation ion patterns for each eluting molecule into a composite mass spectrum^4^. The deconvolution is particularly computationally challenging for complex biological systems where co-elution of compounds is inevitable as raw GC-MS data consist of mass spectra originating from hundreds-to-thousands of molecules.

Annotation of GC-MS data is achieved by matching the deconvoluted fragmentation spectra against reference spectral libraries of known molecules. The 70eV energy for ionizing electrons in GC-MS was set as the standard early, making it possible to use decades-old EI reference spectra for annotation ^5,6^ and compare EI data across instruments. There are now ~1.2 million reference spectra, accumulated and curated over a period of >50 years, that are commercially or publicly available for the annotation of GC-MS data ^6,7^. To date, many analytical tools and several repositories for GC-MS data have been introduced ^5,8–16^. Despite these developments, much GC-MS data processing is restricted to vendor-specific formats and software (e.g. VocBinBase^15^ uses Leco ChromaTOF data). Moreover, the deconvolution requires multiple parameters to be set by the user or manual peak integration. Further, none of the tools are integrated into a mass spectrometry/metabolomics public data repository that retains every setting and result of an analysis job, features that are vital for reproducibility of data processing. A public informatics resource that can not only be integrated with a public repository, but also perform GC-MS deconvolution, alignment, and mass spectral library matching for large numbers (>100) of data files is needed. Technical reasons, such as the lack of a shared and uniform data format, often preclude GC-MS data comparison between different laboratories and prevents taking advantage of repository-scale information and community knowledge about the data. This, coupled to a lack of incentive to deposit data into public domain, leads to GC-MS datasets being infrequently shared and rarely reused across studies and/or biological systems ^15,17–21^.

One of the developments that enabled finding additional structural relationships within mass spectrometry data is spectral alignment, which forms the basis for molecular networking ^22–26^. Here, we develop a repository-scale analysis infrastructure for GC-MS data enable to create networks within the Global Natural Products Social (GNPS) Networking platform. GNPS promotes Findable, Accessible, Interoperable, and Reusable (FAIR) use practices for mass spectrometry data ^27^. The community infrastructure can be accessed at https://gnps.ucsd.edu under the header “GC-MS EI Data Analysis”.

## Results

### Creating a web-based scalable strategy for spectral deconvolution

Current EI spectral deconvolution strategies can save settings and apply them to the next analysis, but require initial manual parameter setting (e.g. AMDIS ^5^, MZmine/ADAP ^8^, MS-DIAL ^9^, PARAFAC2 ^12^); some require extensive computational skills to run (e.g. XCMS ^28^, eRah ^14^). Although batch modes exist, they do not enhance deconvolution quality by utilizing information from other files of the dataset. To use this across-file information, improve scalability of spectral deconvolution, and eliminate manual parameter setting, we developed an algorithmic learning strategy for deconvolution of entire datasets (Figure 1a-f). We deployed this functionality within GNPS/MassIVE^29^ (Figure 1f-i). To promote analysis reproducibility, all GNPS jobs performed are retained in the “My User” space and can be shared as hyperlinks in collaborations or publications.

**Figure 1.**
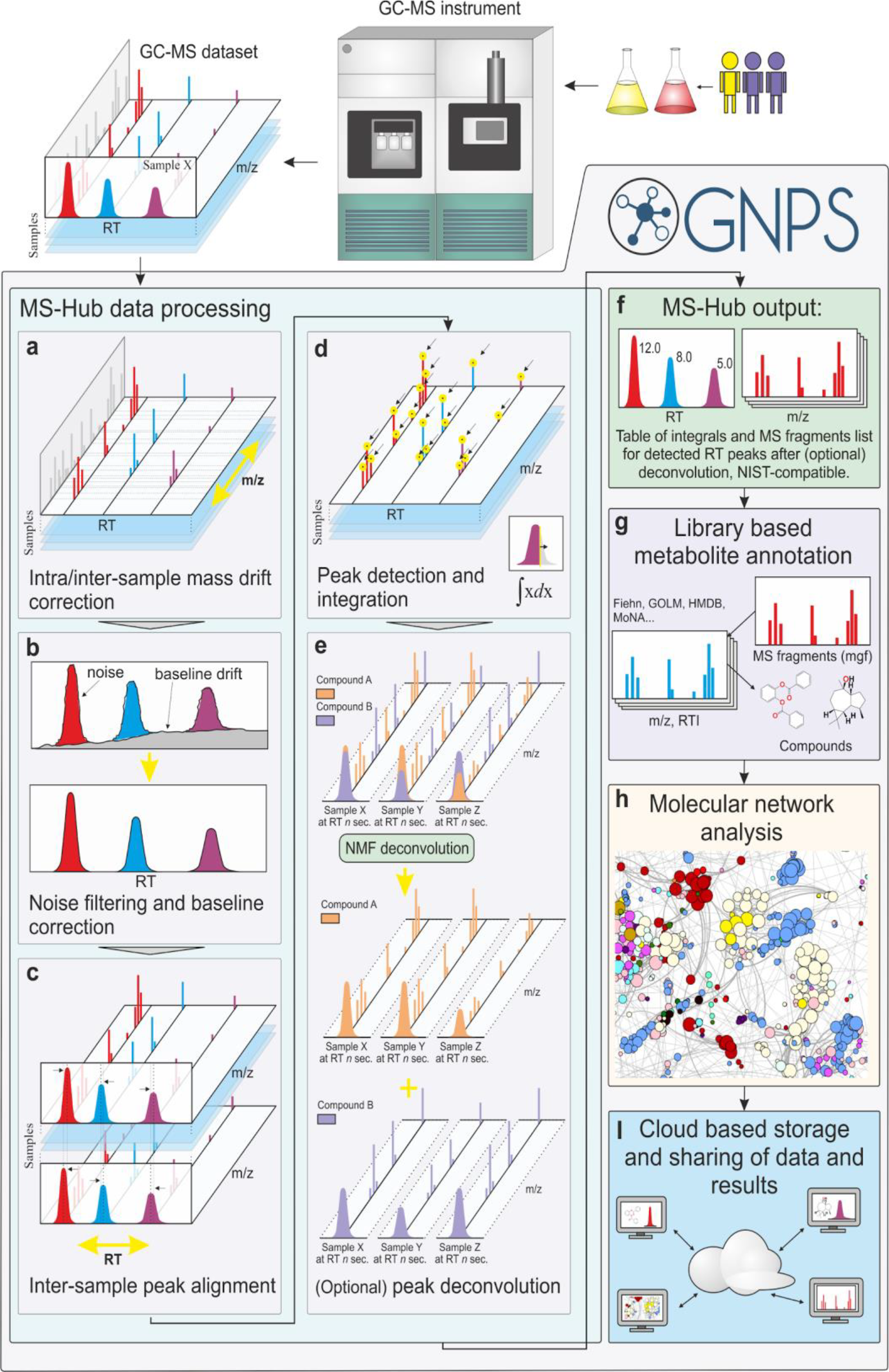
Schematic representation of the MSHub processing pipeline within GNPS. MSHub accepts netCDF, mzML formats of any EI GC-MS data for input. **a**) Spectra are aligned and binned in *m/z* dimension noise is filtered and **b**) baseline is corrected in each spectrum in RT dimension to address issues such as baseline rise with thermal gradient due to bleeds. **c**) Common profile established across entire dataset and peaks in RT dimension are aligned to it using FFT-accelerated correlation in full resolution via iterative approach. **d**) Fast peak detection picks and integrates peaks to generate both peak integrals for all samples and their common fragmentation patterns. For datasets with more than 5 samples peak deconvolution step **e**) is employed to separate overlapping peaks with different patterns across samples using NMF approach. **f**) MSHub produces peak integrals for all samples and canonical fragmentation patterns. **g**) GNPS employs either public or user-provided reference libraries to annotate peaks. **h**) Molecular networks are built for further metabolite analysis. **i**) Data and results are shared between users via GNPS’s cloud architecture. NMF - Non-negative matrix factorization, FFT - Fast Fourier Transform, RT - retention time, *m/z* - mass-to-charge ratio.

Classically, when performing spectral deconvolution of GC-MS data, the user defines parameters specific to their data to the best of their abilities. The user must therefore have a thorough understanding of the characteristics (i.e., peak shape, peak width, resolution etc.) of the particular GC-MS data set before spectral deconvolution. In our approach, the parameters for spectral deconvolution (*m/z* drift of the ions, peak shape - slopes of raising and trailing edges, peak shifts, and noise/intensity threshold) are auto-estimated. This user-independent ‘automatic’ parameter optimization is accomplished via fast Fourier transformation, multiplication, and inverse Fourier transformation for each ion across entire data sets, followed by an unsupervised matrix factorization (one layer neural network): Figure 1a-e. Then, the compositional consistency of spectral patterns, for each spectral feature deconvoluted across the entire data set, can be summarized as a parameter that we termed “balance score”. The balance score (definition is described in the Methods) gives insight into how well the spectral feature is explained across the entire data set: when high, the spectrum is consistent across different samples. Even when a compound is present in only a few samples in the dataset, as long as the spectral patterns are highly conserved across samples (e.g. not contaminated by spurious noise peaks), it would result in a high balance score. Balance score thus allows discarding low-quality spectra that are more likely to be noise, and provides an orthogonal metric to matching scores when searching spectral libraries. We refer to the dataset-based spectral deconvolution tool within the GNPS environment as “MSHub”. MSHub converts raw GC-MS data of any kind (e.g. **Table S1**) into spectral patterns, enabling molecular networking within GNPS.

All MSHub algorithms operate iteratively for enhanced scalability, using high-performance HDF5 technologies saving settings for each analysis step. The Fourier transform step with multiplication dramatically improves MSHub’s efficiency, resulting in deconvolution times that scale linearly rather than exponentially with the number of files (Figure 2a, **S2**). The GNPS GC-MS workflow can process thousands of files in hours (Figure 2a), which is faster than data acquisition, making data processing no longer a bottleneck. We achieved this performance using out-of-core processing, a technique used to process data that are too large to fit in a computer’s main memory (RAM): MSHub uploads files one at a time into the specific RAM module, data are then processed and deleted from memory, iteratively. Figure 2a illustrates the linear dependency between the number of samples processed and the processing time. Because only one sample is stored in memory at any given time, the workflow memory load is constant. Spectral deconvolution scales linearly because each step in the processing pipeline is linear with respect to time (**Figure S2a-f**), taking ~1 min per file (**Figure S1**). The machine learning approaches gain improved performance with increasing amounts of data, which means that increasing dataset size would boost learning each spectral pattern. Indeed, larger volume of analyzed data leads to better scores of spectral matches for the known compounds in derivatized blood serum samples that were spiked with 37 fatty acid methyl esters (FAMEs) and 17 long-chain hydrocarbons (Figure 2b, c). Cosine and balance score can be jointly used as filters for processing the final results (Figure 2d-f). In the analysis of biological samples, similar trends are found as for the reference dataset: the spectral matching scores against the library increase with increasing number of processed files while their distributions become narrower, a reflection that more data leads to better quality of results (Figure 2g, h). When there are more files deconvoluted, MSHub is leveraged to reduce chimeric spectra and discover more real spectral features, which leads to higher quality spectra and a rise in the number of unique annotations with greater match scores (Figure 2i, j). If the user only has a few files (fewer than 10), spectral deconvolution and alignment should be performed using alternative methods (e.g. MZmine^30^, OpenChrom ^28,31^, AMDIS^5^, MZmine/ADAP ^8^, MS-DIA ^9^, BinBase^15^, XCMS ^32^/XCMS Online^28^, MetAlign^10^, SpecAlign^33^, SpectConnect^11^, PARAFAC2^12^, MeltDB^13^, eRah^14^). Using those tools, molecular networking can be performed in the same fashion as for MSHub, as the library search GNPS workflow accepts input from other tools into the GNPS/MassIVE environment. We have further benchmarked the MSHub against XCMS^28^ (MassIVE dataset MSV000084622) and the quantitative results were nearly identical (the calibration curve was within 99.17% correlation with 0.72% STD, Figure 2k,l).

**Figure 2:**
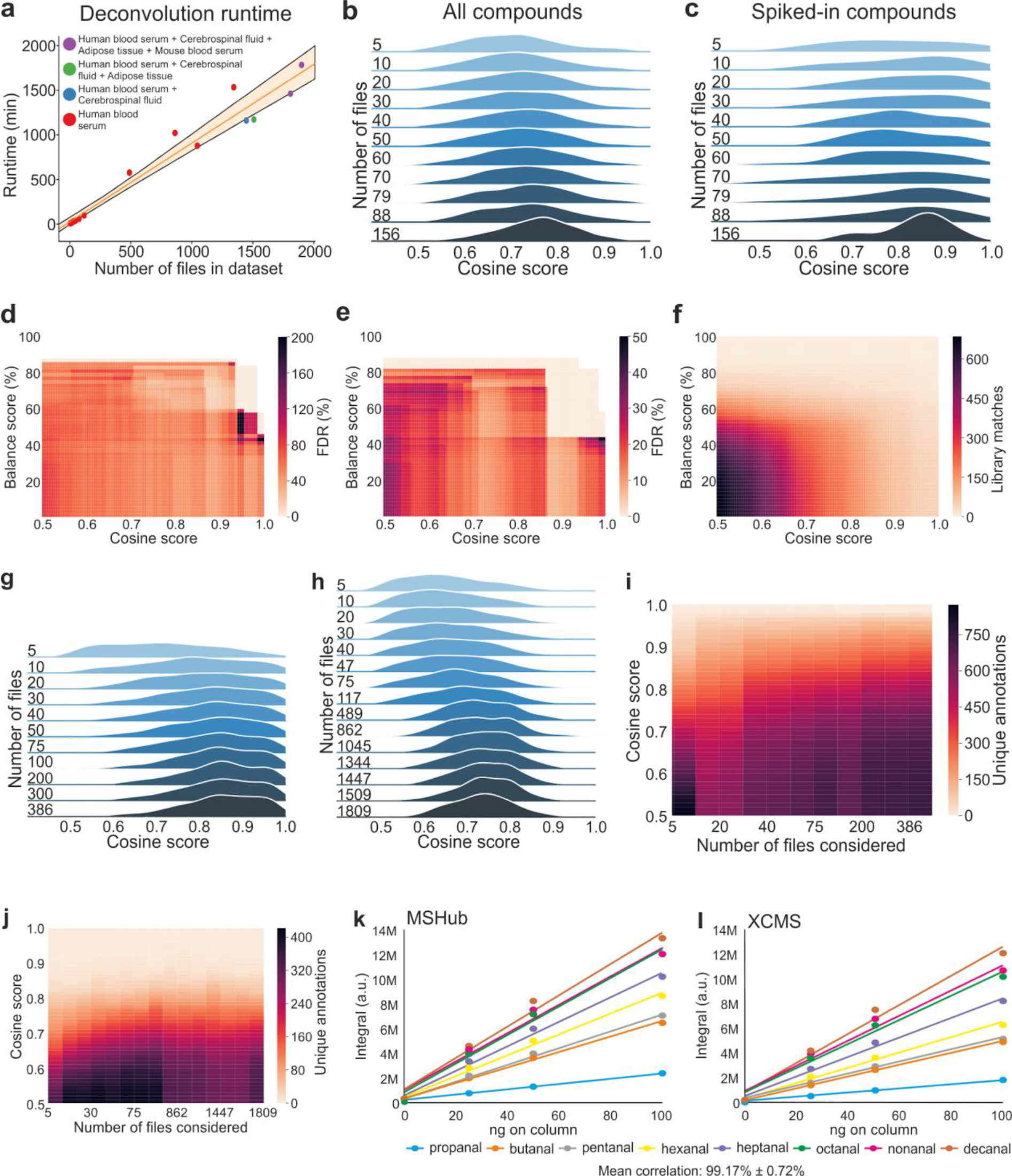
Performance evaluation of dataset-wide deconvolution. **a**) Linear dependence between the number of samples and the processing time on a single compute node. **b**) Distributions of library matching scores for the test dataset of reference compounds spiked in a complex blood serum matrix with an increased volume of input data (**datasets Test1-Test11, Table S1**) for all matches and **c**) for the reference compounds only. **d**) False discovery rate (FDR) for the sub-class^35^ annotations (**dataset Test11 in Table S1**) of the top match and (**e**) top ten matches. More restrictive thresholds minimize misannotations. **f**) Heat map of the number of library matches for spiked compounds. **g**) Distributions of library matching scores for the top match in a study of oesophageal and gastric cancer detection using breath analysis (non-derivatized, **datasets ICL1-ICL11 in Table S1**) and **h**) studies of human and mouse blood serum, adipose tissue and cerebrospinal fluid (silalated, **datasets UCD1-UCD16 in Table S1**). **i**) Heat map of the number of unique annotations (top hit only) for the data across **datasets ICL1-ICL11 in Table S1** and **j**) **datasets UCD1-UCD16 in Table S1**; no balance score filtering applied. Spurious features corresponding to the low cosine tail of the distribution on panels (**g**) and (**h**) are improved as higher volume of the data enhances the frequency domain for deconvolution quality. **k**, **l**) Quantitative integrals of abundances quantitation for the mixture of standards (MassIVE dataset MSV000084622) evaluated using XCMS (**l**) and MSHub (**k**).

Spectral deconvolution using MSHub in GNPS generates an .mgf file that contains deconvoluted spectra with aligned retention times and a feature table of peak areas of features across all files. This generated .mgf spectral deconvolution summary file is used for searching against spectral libraries and for molecular networking. GNPS saves this information, so the deconvolution step does not need to be re-performed for any future analyses. The output results can be downloaded and explored using many external tools,e.g. MetExpert^34^.

### GNPS enables searches against public spectral reference libraries and molecular networking at repository scale

Once the .mgf file is generated by GNPS-MSHub or imported from another deconvolution tool, the spectral features can be searched against public libraries^36^ (currently GNPS has Fiehn^37^, HMDB^38^, MoNA^17^, VocBinBase^15^) or the user’s own private or commercial libraries (such as NIST 2017^39^ and Wiley). Matches are narrowed down based on user-defined filtering criteria such as number of matched ions, Kovats retention index (RI, calculated if hydrocarbon reference values are provided), balance score, cosine score, and abundance. With this release, we also provide additional freely available reference data compiled by co-authors of this manuscript of 19,808 spectra for 19,708 standards. Although the possible candidate annotations can be further narrowed by retention index (RI), they should still be considered level 3, a molecular family, annotation according to the 2007 metabolomics standards initiative (MSI)^40^. Calculation of RIs is enabled and encouraged but not enforced. When multiple annotations can be assigned, GNPS provides all candidate matches within user’s filtering criteria.

No matter how the spectral library is searched in GC-MS, due to the absence of a parent mass, a list of spectral matches is more likely contain mis-annotations, both related (isomers, isobars) or less frequent, entirely unrelated compounds^5^. However to spot misassignments at the molecular family level, we propose to explore deconvoluted GC-MS data via molecular networking, a strategy that has been effective for LC-MS/MS data. In the case of EI, unlike in LC-MS/MS where the precursor ion mass is known, the molecular ion is often absent. For this reason, the molecular networks are created through spectral similarity of the deconvoluted fragmentation spectrum without considering the molecular ion. For GC-MS data that do have a molecular ion or precursor ion mass, e.g. from chemical ionization (CI) or with MS/MS spectra, the feature-based molecular networking workflows should be used^29,41^. We explored clustering patterns for the EI data (**Figure S4**) and observed that the EI-based cosine similarity networks are predominantly driven by structural similarity (**Figure S4a**) ^35^. These EI networks can be used to visualize chemical distributions and guide annotations (**Figure S5**). Networking enables data co- and re-analysis, as it is agnostic to the data origin once the features are deconvoluted. To demonstrate this, we have built a global-network of various public GC-MS datasets deposited on GNPS (38 datasets comprising ~8,500 GC-MS files, Figure 3c). These data encompass various types of samples, modes of sample introduction etc. and thus the global network is a snapshot of all chemistries detectable by GC-MS (Figure 3c-e, **S6**). Prior to networking, we applied a balance score of 65%, which allowed us to remove a bulk of spurious low quality matches (Figure 3 a,b). The balance score filter ensures that the best-explained deconvoluted features are matched against the reference library. The annotation is usually done by ranking potential matches according to a similarity measure (forward match, reverse match, and probability^42,43^) and when possible, filtering by retention index then reporting the top match. Molecular networking can further guide the annotation at the family level by utilizing information from connected nodes (**Figure S5**) rather than focusing on individual annotations^44^. The global network can be colored by metadata such as sample type (Figure 3c), derivatized vs. non-derivatized, instrument type or other metadata (**Figure S6**) to reveal interpretable patterns. When coloring the data by sample type, for example, a cluster of nitrogen-containing heterocyclic compounds was observed to be unique to dart frogs from Dendrobatoidea superfamily (Figure 3e), while the long-chain ketones occur in cheese and beer (**Figure S7**). To highlight the broader utility of GNPS GC-MS and GC-MS based molecular networking, 6 supplemental videos were created that carry the user through how to navigate and perform analysis with the tools (**Supporting Videos 1-6**).

**Figure 3:**
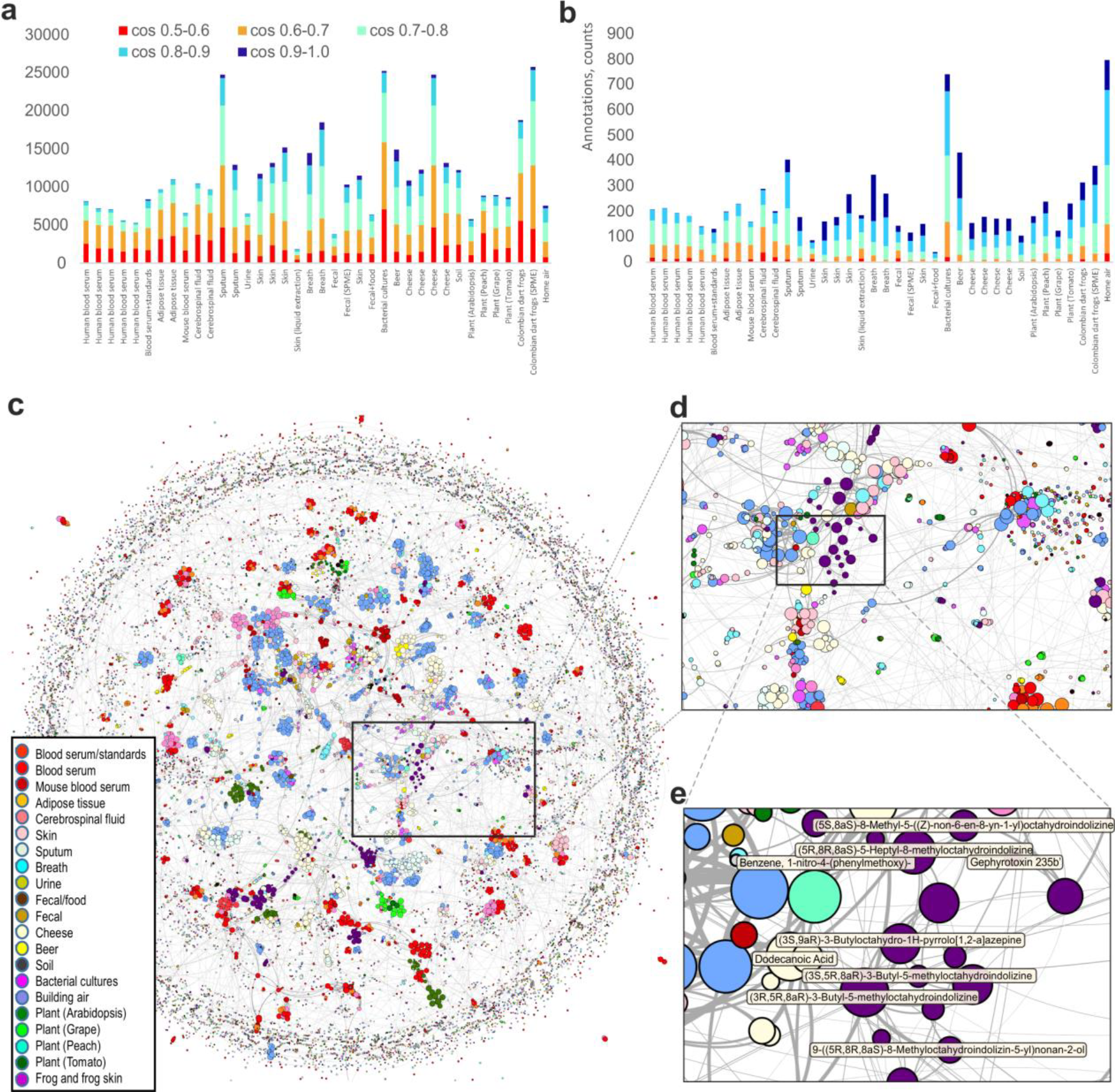
Molecular networking of GC-MS data in GNPS. Features that are annotated (pass library match threshold of 0.5) are included **a)** without filtering and **b)** with a 65% balance score filtering. **c)** Global network containing 35,544 nodes from 8,489 files in 38 GNPS datasets for different types of samples, including human, derived from various animal, plant, microbial and environmental samples. Nodes are connected if cosine > 0.5. The size of the node is proportional to the number of nodes that connect to it ^45^, the edge thickness is proportional to the cosine score (**Figure S3**), the annotation is the top match with cosine above 0.65. d) The inset shows the zoomed-in portion of the network. e) Close up of a cluster of compounds found in the dart frog skin samples with the top spectral library match shown - all nodes are nitrogen heterocyclic alkaloids such as gephyrotoxin ^46^ that are unique to these frogs.

The output from GNPS deconvolution, annotations, and molecular network analysis can be exported for use in a statistical analysis environment such as Qiime ^47,48^, Qiita ^49^, or MetaboAnalyst ^50,51^, or for data visualization in tools such as Cytoscape ^52^ or Gephi ^53^ (e.g.**Supplementary Figures S4-S7**, Figure 3, 4e-g), or for molecular cartography in ‘ili ^54^ (Figure 4 a-d). To demonstrate how to use GNPS GC-MS for the latter, we collected samples from 52 body locations from one person using a sampling patch that absorbs volatiles (Figure 4 a-d). These samples were subjected to headspace desorption followed by GC-MS, deconvoluted and annotated using the GNPS GC-MS pipeline. The abundances from the deconvoluted spectra are superimposed onto a 3D model of a human (Figure 4 a-d). Using balance filters at 50% and >0.9 cosine, we arrived at annotations that, once visualized, revealed the distributions of skin volatiles. For example, squalene was found on all locations, but less on the feet. Hexanoic acid was most abundant on the chest and armpits. Globulol, an ingredient of the personal care product this individual used on the chest, was most intense on the chest, while phenylenedibenzoate, a skincare ingredient, was found on the face and hands.

**Figure 4.**
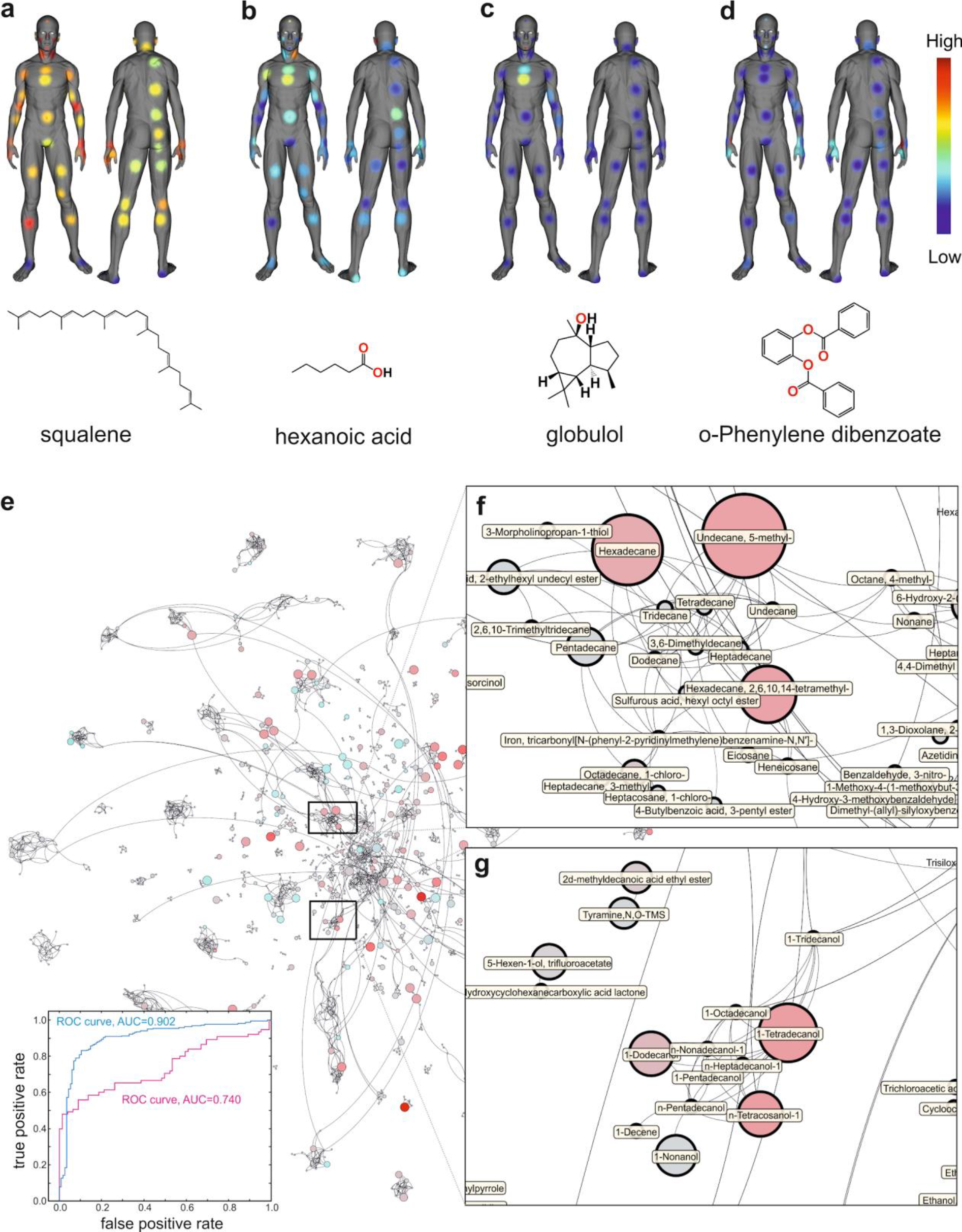
Examples of results with GNPS processed GC-MS data. 3D visualization of human surface volatilome visualized with ‘ili ^54^ as described in the tutorial (https://ccms-ucsd.github.io/GNPSDocumentation/gcanalysis/). Molecular distributions on skin of a volunteer shown for: **a)** squalene, a key component of natural skin grease. Low abundance of squalene is aligned with areas of dry skin ^72^ **b)** hexanoic acid, one of the malodour molecules responsible for the unpleasant sour body odor ^73^ **c)** globulol, naturally occurring in plant essential oils, likely introduced via use of skin cosmetics **d)** phenylenedibenzoate, also introduced via use of a skin product. **e)** Chemical distributions that relate to cancer status are visualized via molecular network that combines two studies. Each node represents a unique mass spectrometry feature obtained from deconvolution. The top annotation is given for matches with cos>0.65. The size of the node represents the importance of the feature for discrimination by the maximum margin criterion ^74^ with leave patient out cross validation of the OGC group vs control volunteers; the color of the node represents average fold-change in abundance between OGC vs. control groups (red - higher in OGC, teal - higher in control, gray - neither), the size represents -log(p value), larger circle corresponds to greater values. The inset shows ROC for both studies (Study 1 - blue, Study 2-red). **f)** Example of cluster of hydrocarbons and **g)** long-chain alcohols. Both human studies are approved by the institutional review boards as described in the Methods.

We also conducted two studies (Study 1: n=631 samples and Study 2: n=219 samples, respectively) on breath analysis associated with oesophageal and gastric cancers (OGC, Figure 4 e-g). In breath, biological signatures are usually obscured by intra- and inter-subject variability, experimental conditions, e.g. ambient air quality, different diets etc.Biologically relevant compounds are often present at low abundances. Both studies predicted OGC (inset in Figure 4e). The next important step is to consider features that are the most discriminant between categories of interest (OGC vs. control) to investigate whether their chemical identity can be linked to a plausible biological rationale. However, even though OGC prediction was achieved in each study, the “OGC signatures” do not appear to overlap between the two studies, which is very typical for breath analysis field in general^55–58^. As molecular networking organizes chemically similar compounds into clusters, it facilitates recognition of patterns at a chemical family level. Exploring the two studies as a single network revealed an increase of related but not identical medium/long chain alcohols, aldehydes and hydrocarbons (Figure 4 e-g). Only a handful of these compounds appeared as top discriminating features in either study. Aldehydes are known to be found endogenously, mostly due to lipid peroxidation, and have been proposed as potential biomarkers in exhaled breath in several different types of cancer including lung ^59–62^, breast^63^, ovarian^64^, colorectal^65^, and, most notably, OGC ^66–68^. The alkanes and methyl branched alkanes have not been previously associated with oesophageal or gastric cancer, but have been associated with lung and breast cancer in exhaled breath ^60,61,63,69,70^. Lipid peroxidation of polyunsaturated fatty acids in cell membranes generates alkanes that can then be excreted in the breath^71^, which makes their observation in relation to OGC biologically plausible. Although few individual alkanes were found significantly increased in OGC cohort in both studies, none of them overlapped, and without considering these data as a single network, association of long-chain alkanes with OGC would be far more difficult to recognize.

## Discussion

GNPS provides a platform for data sharing and accumulation of public knowledge. Community adoption of GNPS has sharply increased the volume of MS data in the public domain^7^. It has also spurred new tools development (MASST^75^, FBMN^41^, ReDU^76^) and enabled many biological discoveries. Due to the fundamental differences between CID and EI fragmentation, the GNPS infrastructure could not previously support the analysis of EI data. Adopting existing solutions for deconvolution was not possible as all of them required too much manual input from the user and could not operate at repository scale. Here we used an unsupervised non-negative matrix factorization and a Fast Fourier Transform-based approach to scale the deconvolution step. Such strategies are most effective when large-scale datasets become available, as features can be extracted with increasing quality of fragmentation patterns, as defined by the balance score. Currently, all 1D EI GC-MS data are amenable and we will extend the same approach to 2D GC-MS data.

These features can then be subjected to EI-based molecular networking. The algorithm for molecular networking within GNPS had to be modified to accommodate EI data to function without molecular ion information and can reinforce candidate annotations to level 3 by assessing if the annotations are similar at the family level and if annotations share chemical class terms. Such analysis can now be achieved with the data at repository scale, enabling co- and re-analysis of GC-MS data. Here we show how the co-analysis could be beneficial for two cancer breathomics data sets, but in the same fashion other breathomics (or other volatilome) data can now be co-analyzed with these datasets as long as they are publicly available in an open file format. Co-analyzing multiple disparate GC-MS studies would be challenging otherwise. Further, when considering GC-MS data as networks, in addition to conventional statistical approaches, strategies such as networks on graphs^77^ could be deployed to investigate global biochemical patterns rather than differences in individual compounds. The networks, in principle, are not limited to any one kind of data and can be extended to any number or type of datasets as shown in Figure 3c.

Surprisingly, although GC-MS is the oldest and most established of MS-based methods, and the sheer volume of existing EI reference data accumulated over decades (far exceeding that for any other kind of MS), researchers still use decades-old data analysis strategies. We anticipate that the new GNPS community infrastructure will incentivize moving raw EI data into the public domain for data reuse, comparable to the trajectory for tandem MS^7,75,76^. GC-MS analysis within GNPS/MassIVE will lower the expertise threshold required for analysis, encourage FAIR practices^27^ through reusable deposition of the data in the public domain, and promote data analysis reproducibility and “recycling” of GC-MS data. Finally, this work is a piece of the puzzle to democratize scientific analysis from all over the world. GC-MS the most widely used MS method, in part due to its competitive operational cost. It is often the only mass spectrometry method available at smaller, e.g. undergraduate institutions, non-metabolomics laboratories, or local testing facilities. The proposed infrastructure will enable labs with fewer resources, including those from developing countries, to have free access to data and reference data in a uniform format, and to free, powerful computing infrastructures.

## Data and code availability

All of the data used in preparation of this manuscript are publicly available at the MassIVE repository at the UCSD Center for Computational Mass Spectrometry website (https://massive.ucsd.edu). The dataset accession numbers are: #1 (MSV000084033), #2 (MSV000084033), #3 (MSV000084034), #4 (MSV000084036), #5 (MSV000084032), #6 (MSV000084038), #7 (MSV000084042), #8 (MSV000084039), #9 (MSV000084040), #10 (MSV000084037), #11 (MSV000084211), #12 (MSV000083598), #13 (MSV000080892), #14 (MSV000080892), #15 (MSV000080892), #16 (MSV000084337), #17 (MSV000083658), #18 (MSV000083743), #19 (MSV000084226), #20 (MSV000083859), #21 (MSV000083294), #22 (MSV000084349), #23 (MSV000081340), #24 (MSV000084348), #25 (MSV000084378), #26 (MSV000084338), #27 (MSV000084339), #28 (MSV000081161), #29 (MSV000084350), #30 (MSV000084377), #31 (MSV000084145), #32 (MSV000084144), #33 (MSV000084146), #34 (MSV000084379), #35 (MSV000084380), #36 (MSV000084276), #37 (MSV000084277), #38 (MSV000084212).

All of the GNPS analysis jobs for all of the studies are summarized in **Table S1**.

The source code of the MSHub software is available online at Github (version used in GNPS) (https://github.com/CCMS-UCSD/GNPS_Workflows/tree/master/mshub-gc/tools/mshub-gc/proc) and at BitBucket (standalone version in MSHub developers’ repository: https://bitbucket.org/iAnalytica/mshub_process/src/master/). Scripts used to parse, filter, organize data and generate the plots in the manuscript are available online at Github (https://github.com/bittremieux/GNPS_GC_fig). Script for merging individual .mgf files into a single file for creating global network is available at Github: https://github.com/bittremieux/GNPS_GC/blob/master/src/merge_mgf.py)The 3D model, feature table with coordinates used for the mapping and snapshots shown on the Figure 4a-d are available at: https://github.com/aaksenov1/Human-volatilome-3D-mapping-

## Methods

These are provided as supporting information. The tools are accessible through gnps.ucsd.edu and the documentation on how to use the GNPS GC-MS Deconvolution workflow and molecular networking workflows can be found here https://ccms-ucsd.github.io/GNPSDocumentation/gcanalysis/. Representative examples and short “how to” video can be found here: https://www.youtube.com/watch?v=yrru-5nrsdk&feature=youtu.be

https://www.youtube.com/watch?v=MblruOSglgI&feature=youtu.be

https://www.youtube.com/watch?v=iX03r_mGi2Q&feature=youtu.be

https://www.youtube.com/watch?v=mv-fw2zSgss&feature=youtu.be

https://www.youtube.com/watch?v=nUhCZ9LwoM4&feature=youtu.be

https://www.youtube.com/watch?v=_PehOiBqzzY&feature=youtu.be

## Author contributions

PCD, AAA, MW, LFN came up with the concept of GNPS for GC-MS data.

KV designed and supervised MSHub platform development

IL, DV, VV, KV developed the MSHub platform

MW, ZZ, AAA developed the workflows

AAA, ZZ, MW, BBM, RSB performed infrastructure testing and benchmarking

AAA, ZZ assessed EI-based molecular networking

WB generated plots for MSHub algorithm performance testing

ZZ, AA, ME generated molecular network plots

ME, JJJvdH adapted the MolNetEnhancer workflow for GC-MS Molecular Networks

AAA, AVM, MP, KJ, KD conducted 3D skin volatilome mapping studies

SD, IB, GH conducted oesophageal and gastric breath analysis cancers detection study

AAA, ZZ, MP, MW converted and added public libraries to GNPS

AAA, AVM, SD, BBM, MG, CC, AA, JM, RQ, AB, AAO, DP, AMS, SPC, TOM, MCB, CDN, EZ, VA, EHF, RG, MMM, IM, SE, PLB, BA, RL, YG, SP, AP, GD, BLB, AF, NS, KG, CS, RC, MG, JM, JUS, DB, SA, AF generated GC-MS data

RSB, LNK, AAA assembled the initial version of the public reference spectra library

RS created MZmine export module for GNPS GC-MS input files and RI markers file export

AAA, RS, IB, AAO, AMS, BA, MG, KNM, RSB produced training videos

AAA, MNE, MG, LFN wrote and compiled tutorials and documentation

PCD, AAA, WB, KV, RM, RK wrote the paper

## Ethics/COI declaration

Pieter C. Dorrestein is a scientific advisor for Sirenas LLC.

Mingxun Wang is a consultant for Sirenas LLC and the founder of Ometa labs LLC.

Alexander A. Aksenov is a consultant for Ometa labs LLC.

<u>**Online Tutorial:** https://ccms-ucsd.github.io/GNPSDocumentation/gcanalysis/</u>

## Acknowledgments

The conversion of the data from different repositories was supported by R03 CA211211 on reuse of metabolomics data, to build enabling chemical analysis tools for the ocean symbiosis program, the development of a user-friendly interface for GC-MS analysis was supported by the Gordon and Betty Moore Foundation through Grant GBMF7622. The UC San Diego Center for Microbiome Innovation supported the campus wide SEED grant awards for data collection that enabled the development of some of this infrastructure. PCD was supported by NSF grant IOS-1656475. KV and IL are very grateful for the support of Vodafone Foundation as part of the project DRUGS/DreamLab. AB was supported by the National Institute of Justice Award 2015-DN-BX-K047. Additional support for data acquisition and data storage was provided by P41 GM103484 Center for Computational Mass Spectrometry, the collection of data from the HomeChem project was supported by the Sloan Foundation. GBH, SD, IL, KV and IB are grateful for the support of the OG cancer breath analysis study by the NIHR London Invitro Diagnostic Co-operative and Imperial Biomedical Research Centre, Rosetrees and Stonegate Trusts and Imperial College Charity. IB acknowledges the contribution of Qing Wen and Dr Michelangelo Colavita for the production of the training video. CC was supported by the Research Foundation Flanders (FWO), with support from the industrial research fund of Ghent University. WB was supported by the Research Foundation Flanders (FWO). AAO acknowledges the support of Fulbright Commission and Consejo Nacional de Investigaciones Científicas y Técnicas (CONICET-Argentina). The work of RL and PLB on the dataset 30 was supported by the Metaboscope, part of the "Platform 3A" funded by the European Regional Development Fund, the French Ministry of Research, Higher Education and Innovation, the region Provence-Alpes-Côte d’Azur, the Departmental Council of Vaucluse and the Urban Community of Avignon. SA and ARF acknowledge the PlantaSYST project by the European Unions Horizon 2020 research and innovation programme (SGA-CSA No 664621 and No 739582 under FPA No. 664620). VV acknowledges the support by the National Institute On Alcohol Abuse and Alcoholism award R24AA022057. MG and RC acknowledge the support of the Krupp Endowed Fund grant. A portion of mass spectra in the public reference library was produced within the framework of the State Task for the Topchiev Institute of Petrochemical Synthesis RAS and with the support of the RUDN University Program 5-100. RSB acknowledges support of the State Task for the Topchiev Institute of Petrochemical Synthesis RAS. LNK acknowledges support of the RUDN University Program 5-100. IM acknowledges support of the Israel Science Foundation project number 1947/19 and European Research Council under the European Union’s Horizon 2020 research and innovation program (project number 640384). JS has been supported by NIH/NIAMS R03AR072182, The Colton Center for Autoimmunity, Rheumatology Research Foundation, The Riley Family Foundation and The Snyder Family Foundation. JM acknowledges support from 2017 Group for Research and Assessment of Psoriasis and Psoriatic Arthritis (GRAPPA) Pilot Research Grant and NIH/NIAMS T32AR069515. RG is grateful to the Azrieli Foundation for the award of an Azrieli Fellowship. JJJvdH acknowledges support from an ASDI eScience grant, ASDI.2017.030, from the Netherlands eScience Center-NLeSC. BA was supported by the National Science Foundation (NSF) through the Graduate Research Fellowship Program. GC-MS analyses for collection of the dataset MSV000083743 were supported by the Pacific Northwest National Laboratory, Laboratory Directed Research and Development Program, and were contributed by the Microbiomes in Transition Initiative; data were collected in the Environmental Molecular Sciences Laboratory, a national scientific user facility sponsored by the Department of Energy (DOE) Office of Biological and Environmental Research and located at Pacific Northwest National Laboratory (PNNL). PNNL is operated by Battelle Memorial Institute for the DOE under contract DEAC05-76RLO1830. Authors are grateful to Drs. Marina Vance and Delphine Farmer who have organized the sampling for HomeChem indoor chemistry project (https://indoorchem.org/projects/homechem/) that allowed to collect samples for the dataset MSV000083598. Brandon Ross has assisted with collecting data for the dataset MSV000084348. GC-MS analyses for collection of the datasets MSV000084211 and MSV000084212 were supported by the announcement N757 Doctorados Nacionales and project EXT-2016-69-1713 from Departamento Administrativo de Ciencia, Tecnología e Innovación (COLCIENCIAS), the seed project INV-2019-67-1747 and FAPA project of Chiara Carazzone from the Faculty of Science at Universidad de los Andes, and the grant No. FP80740-064-2016 of COLCIENCIAS. Authors are grateful to Lida M. Garzón, Pablo Palacios, Marco Gonzalez and Jack Hernandez for their contributions collecting the samples, and to Jhony Oswaldo Turizo for designing and manufacturing the amphibian electrical stimulator.

