## Supplementary figures and images for "Algorithmic Learning for Auto-deconvolution of GC-MS Data to Enable Molecular Networking within GNPS"

### Supplemental Figure 1

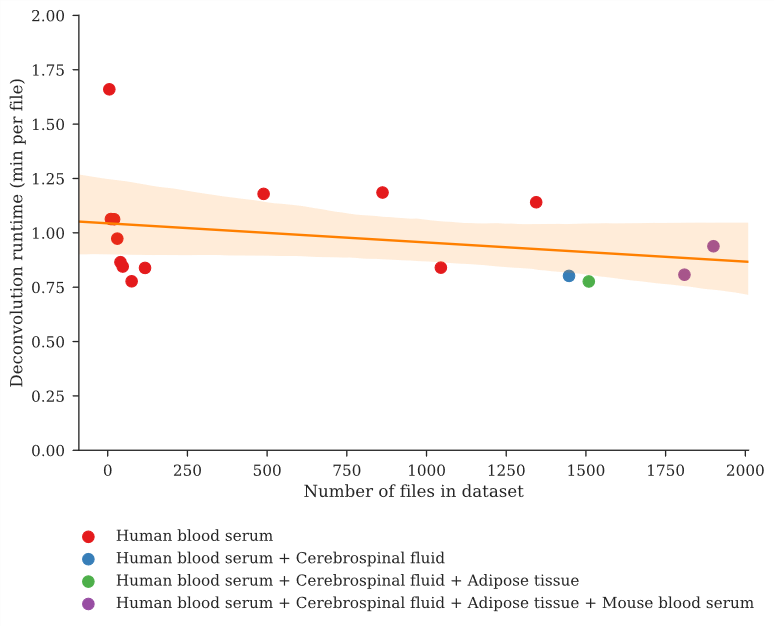

### Supplemental Figure 2

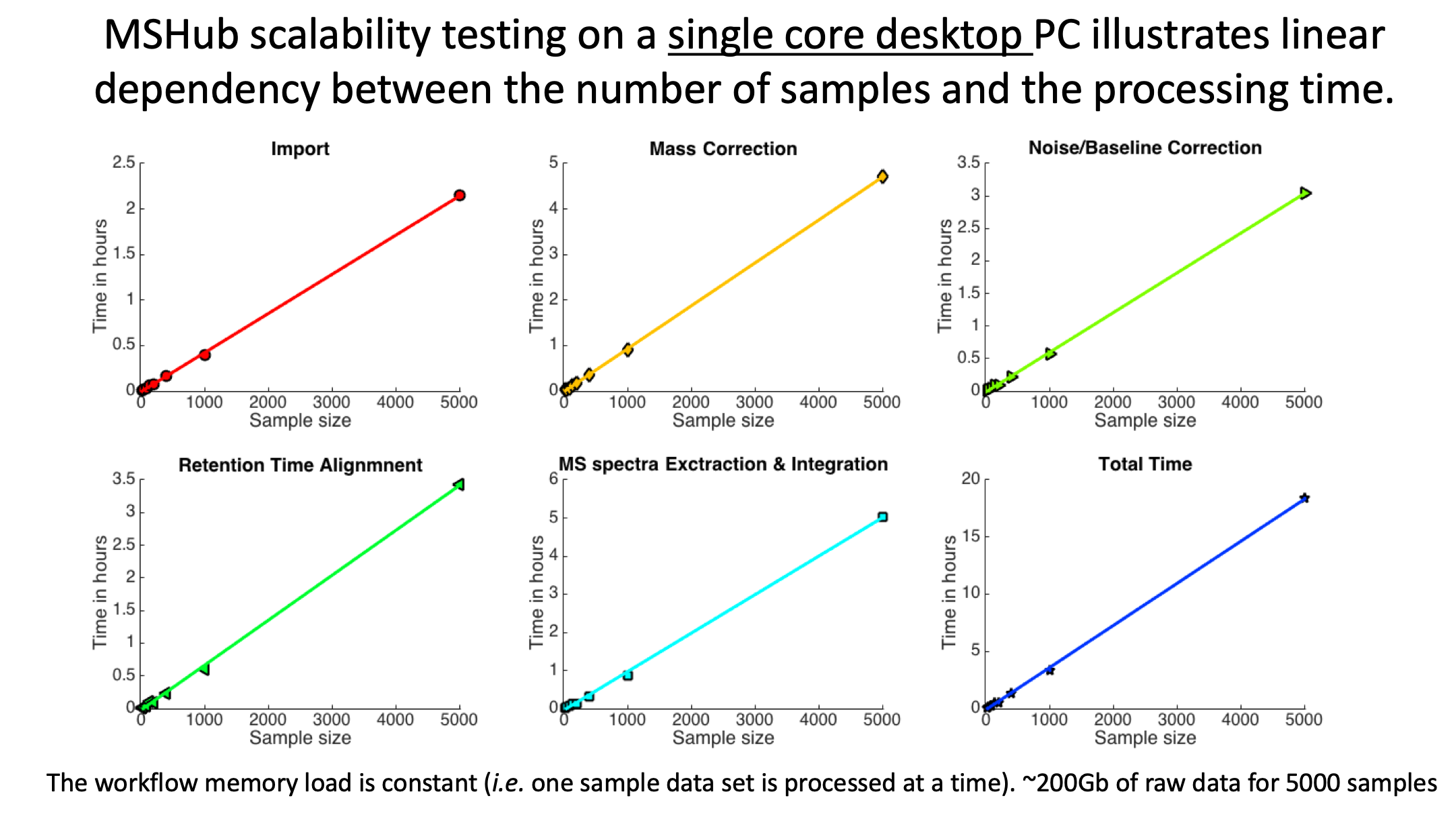

### Supplemental Figure 3

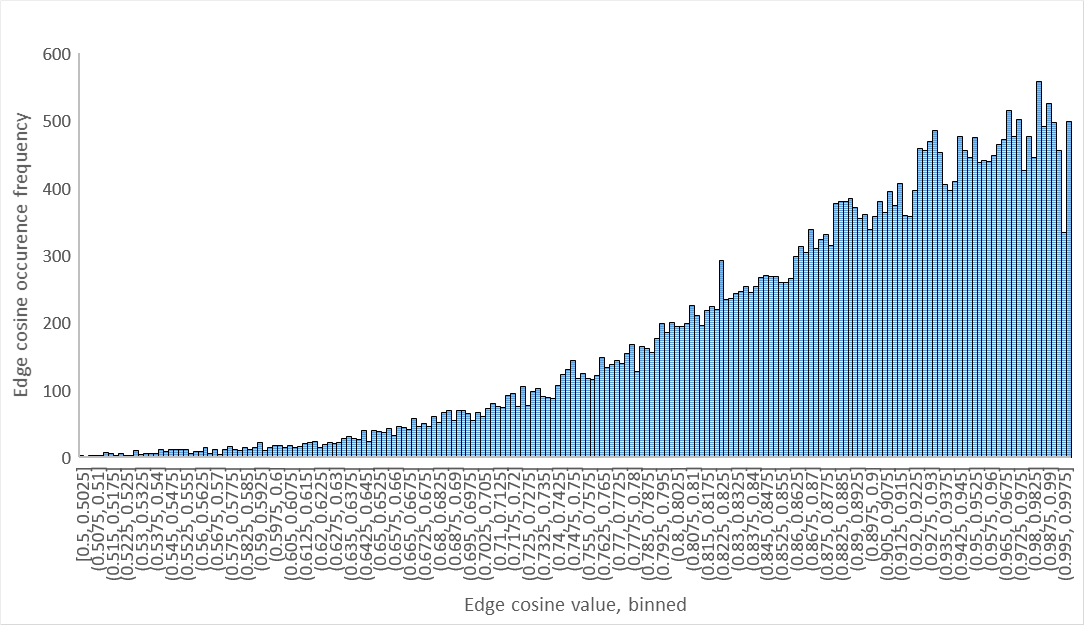

### Supplemental Figure 4a

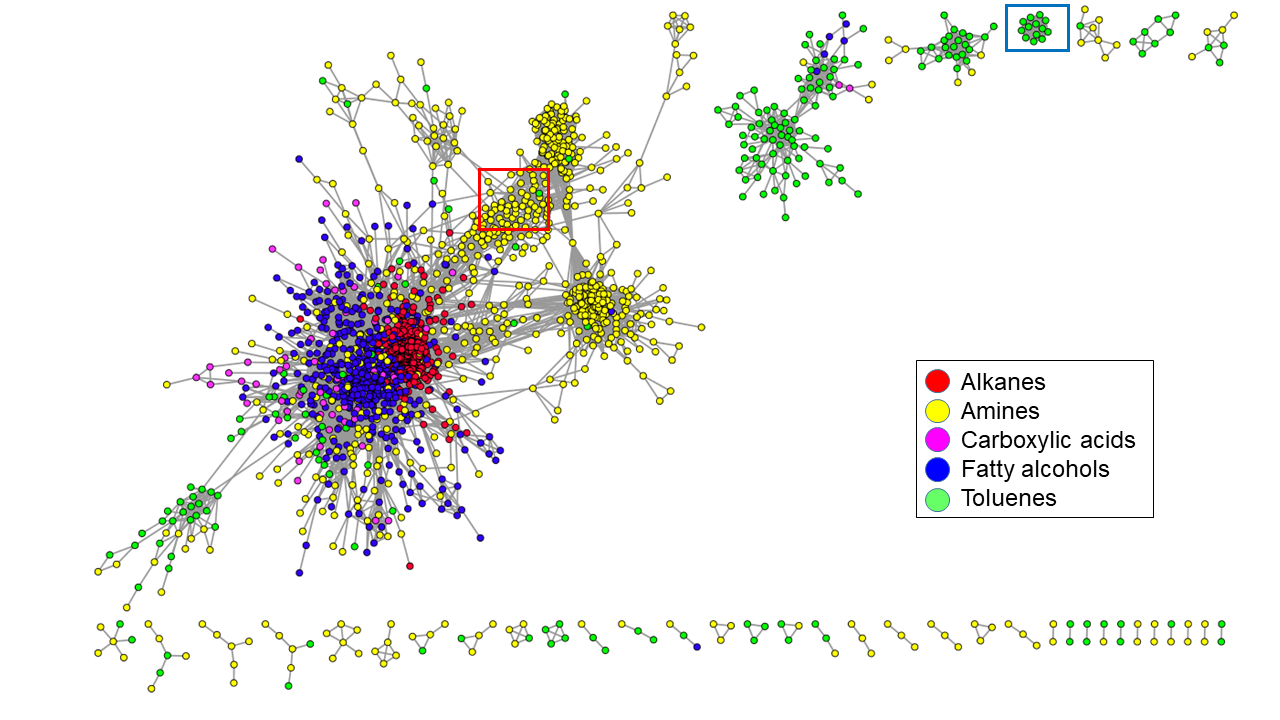

### Supplemental Figure 4b

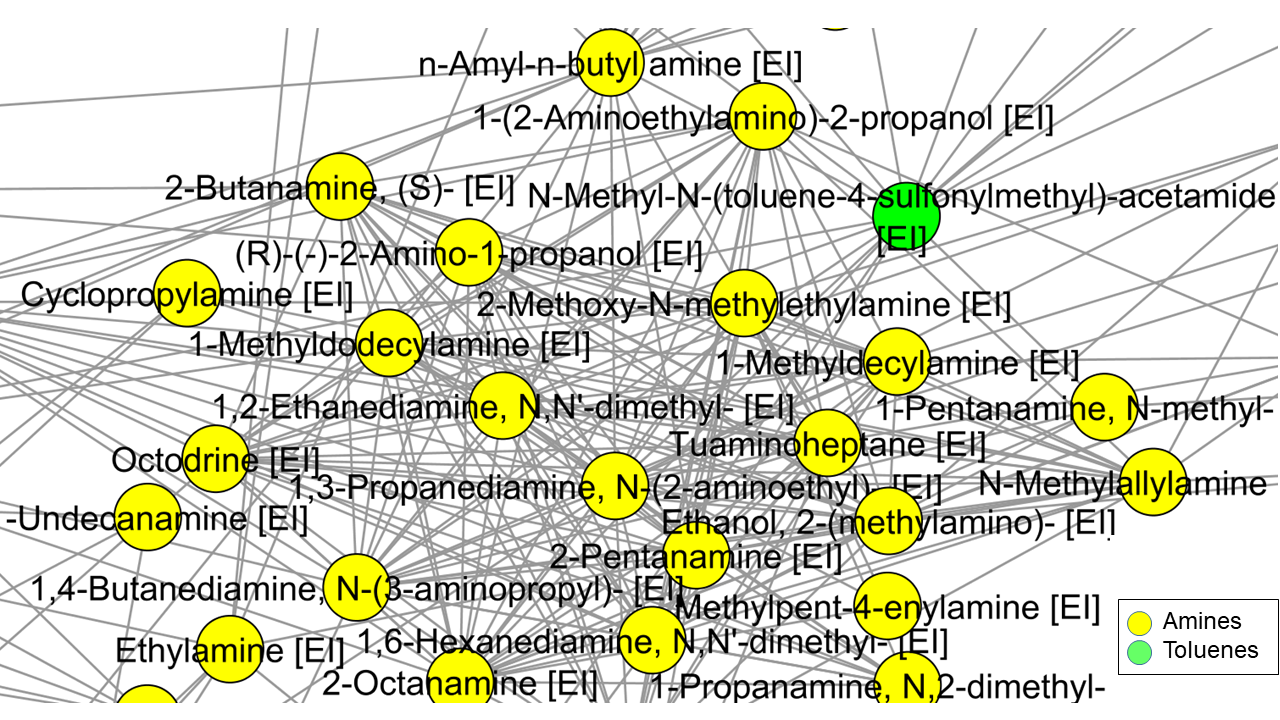

### Supplemental Figure 4c

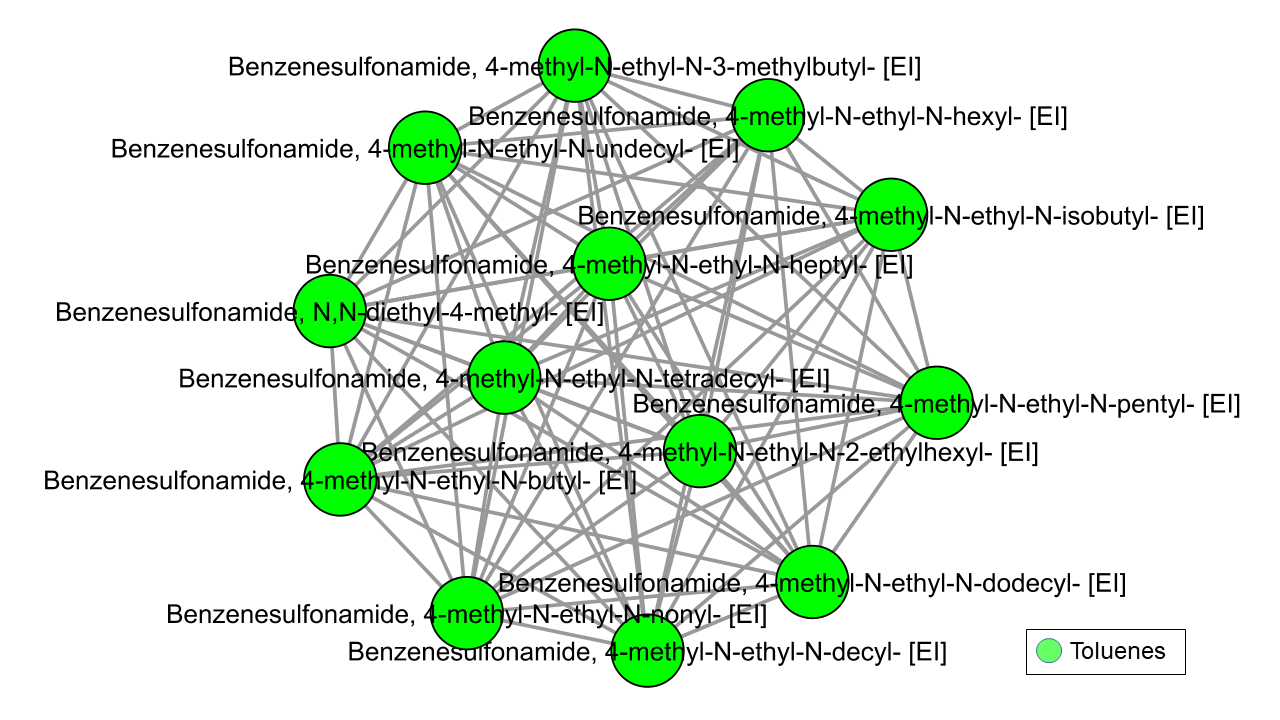

### Supplemental Figure 5a

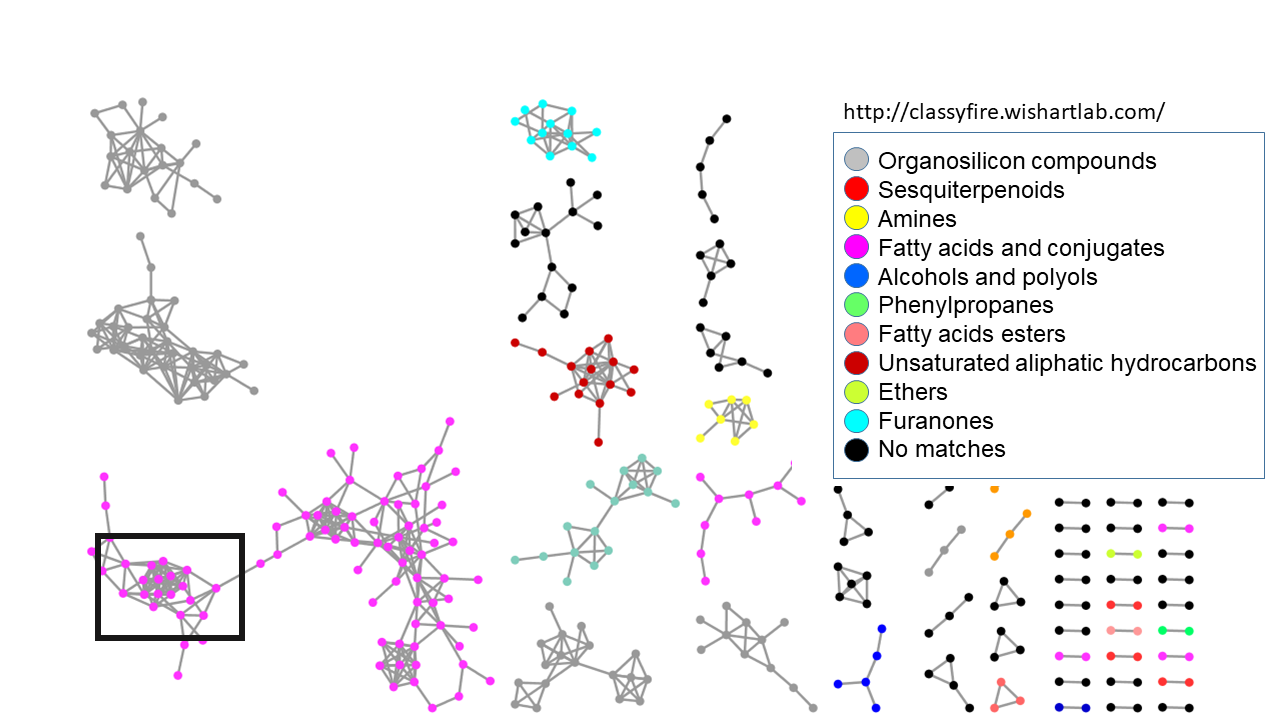

### Supplemental Figure 5b

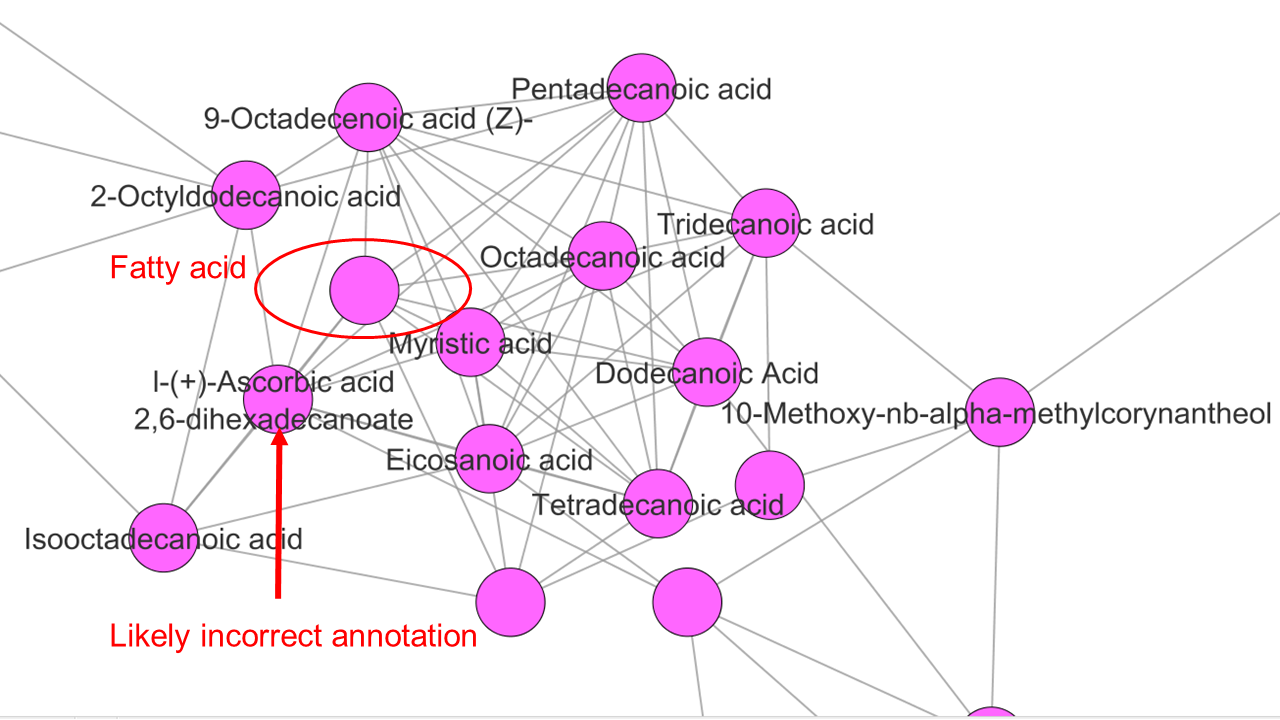

### Supplemental Figure 6a

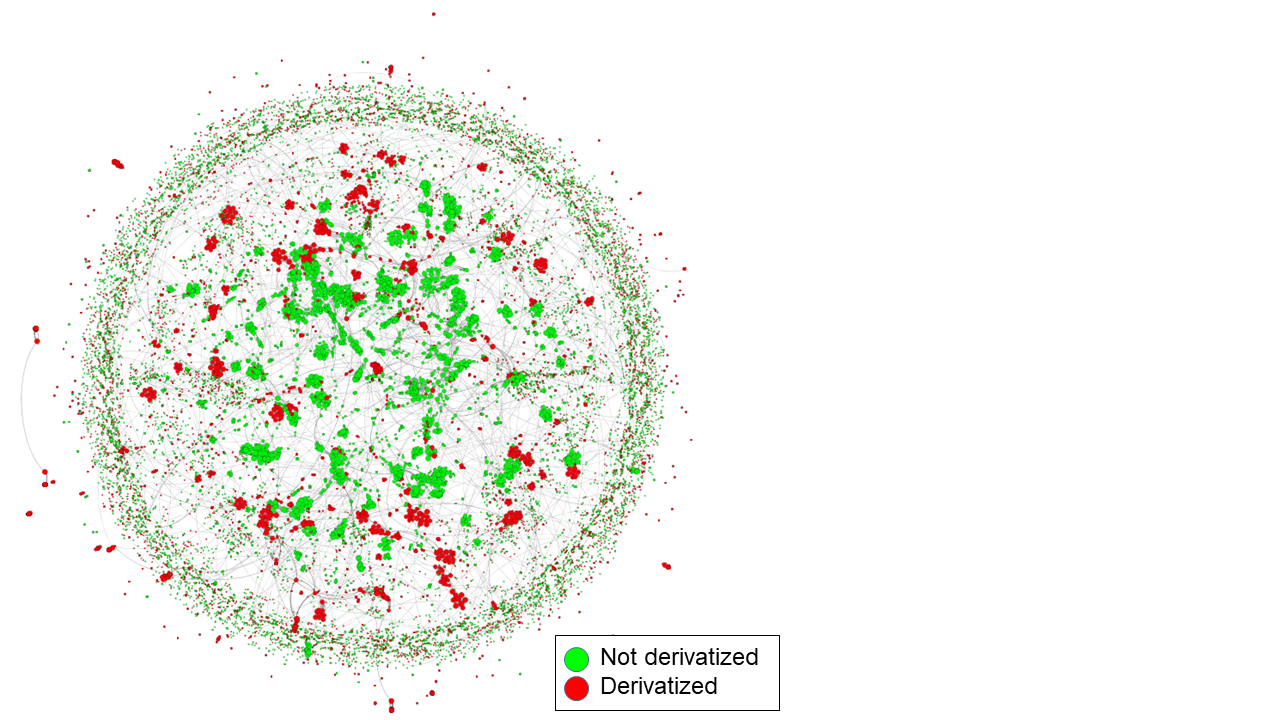

### Supplemental Figure 6b

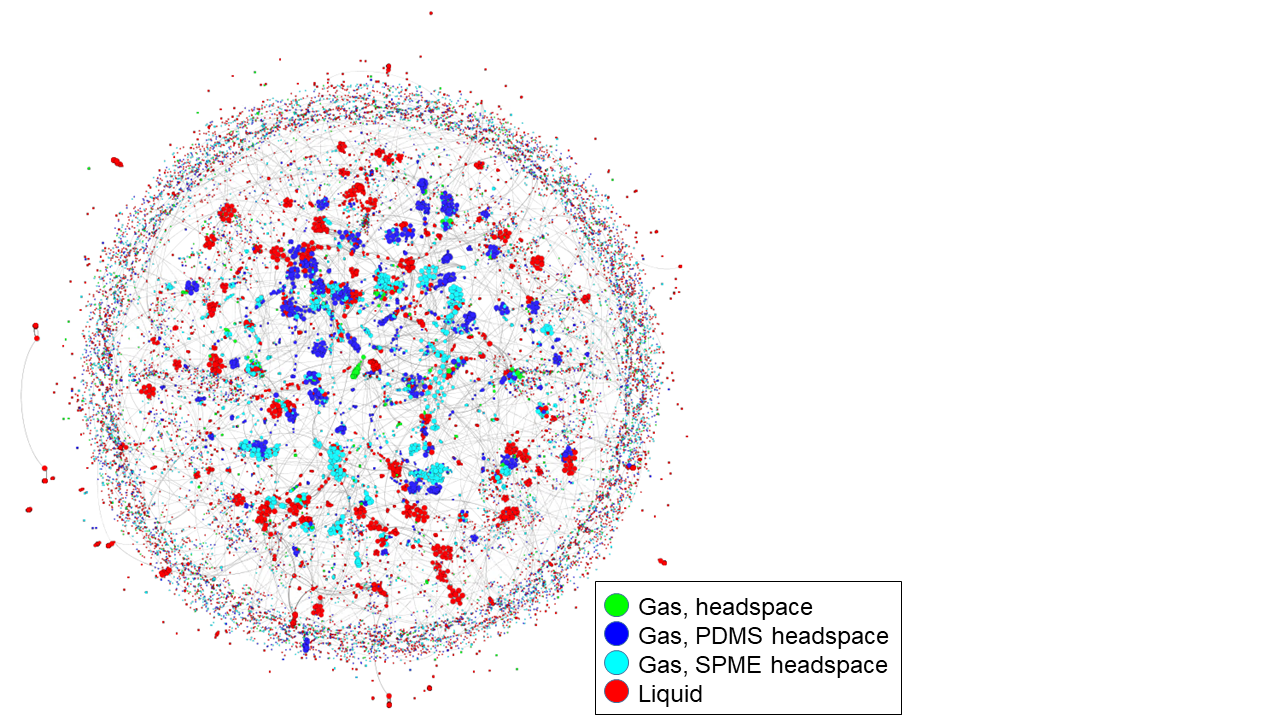

### Supplemental Figure 6c

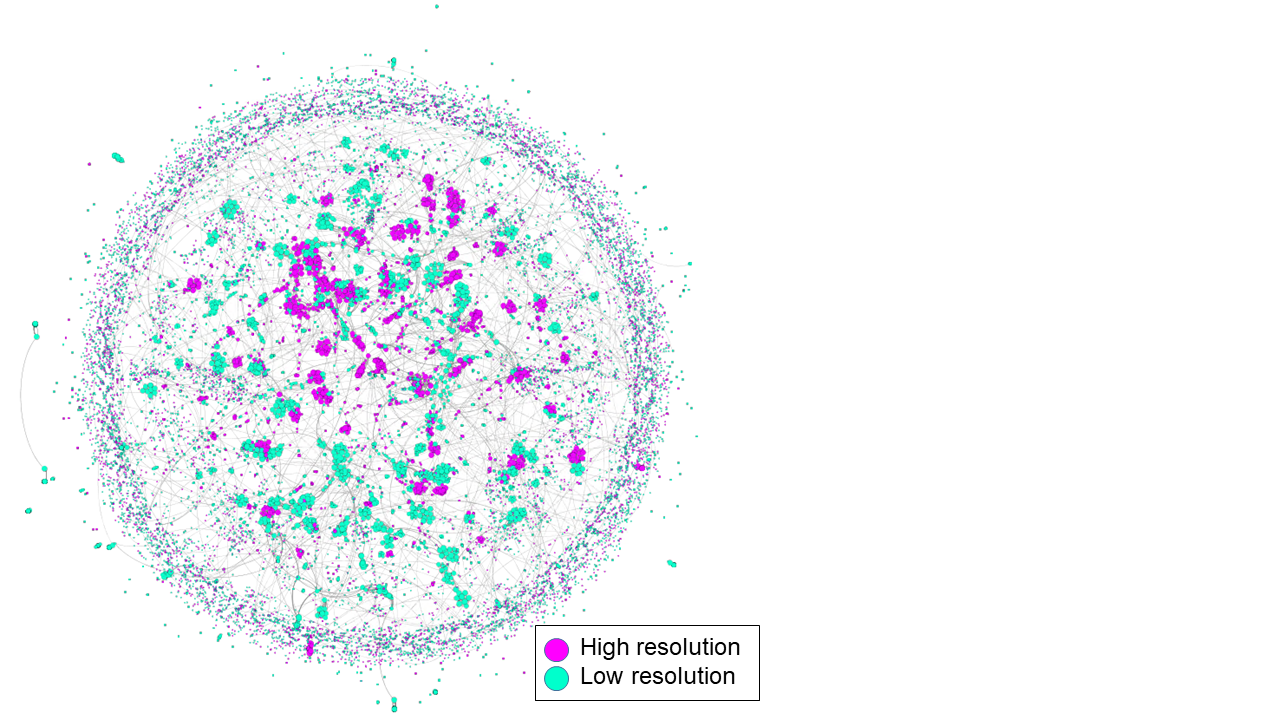

### Supplemental Figure 7a

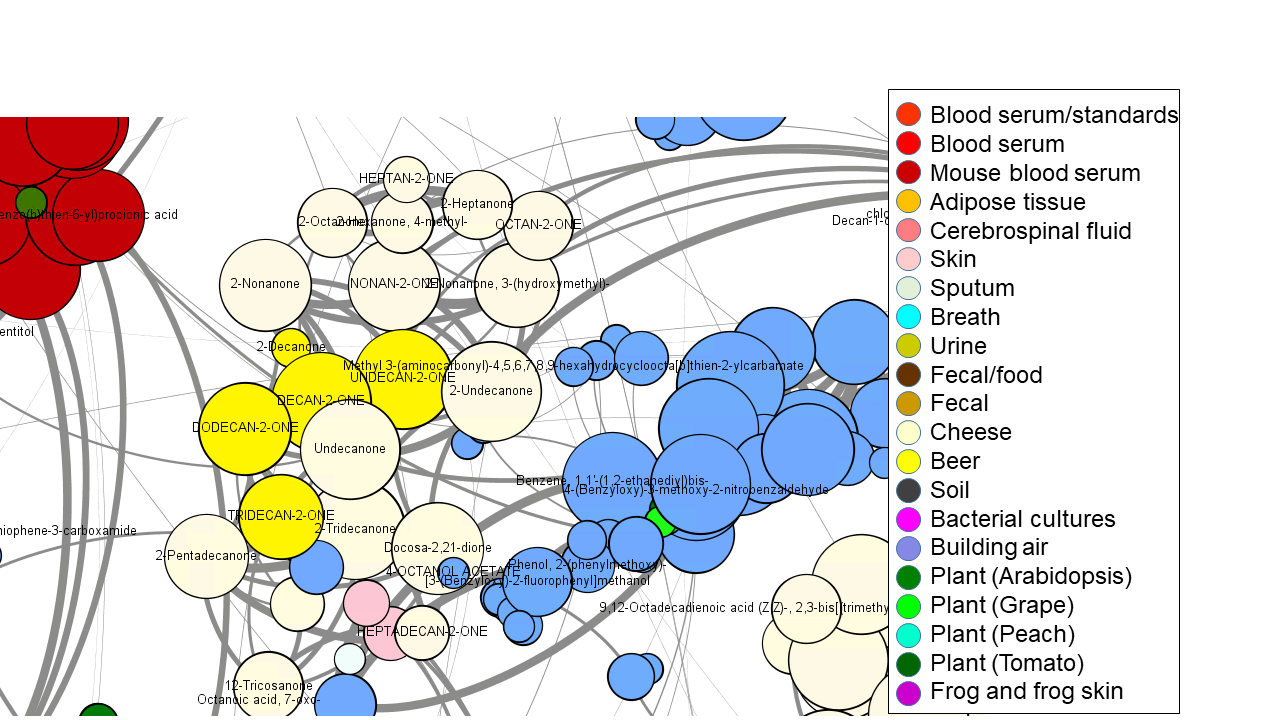

### Supplemental Figure 7b

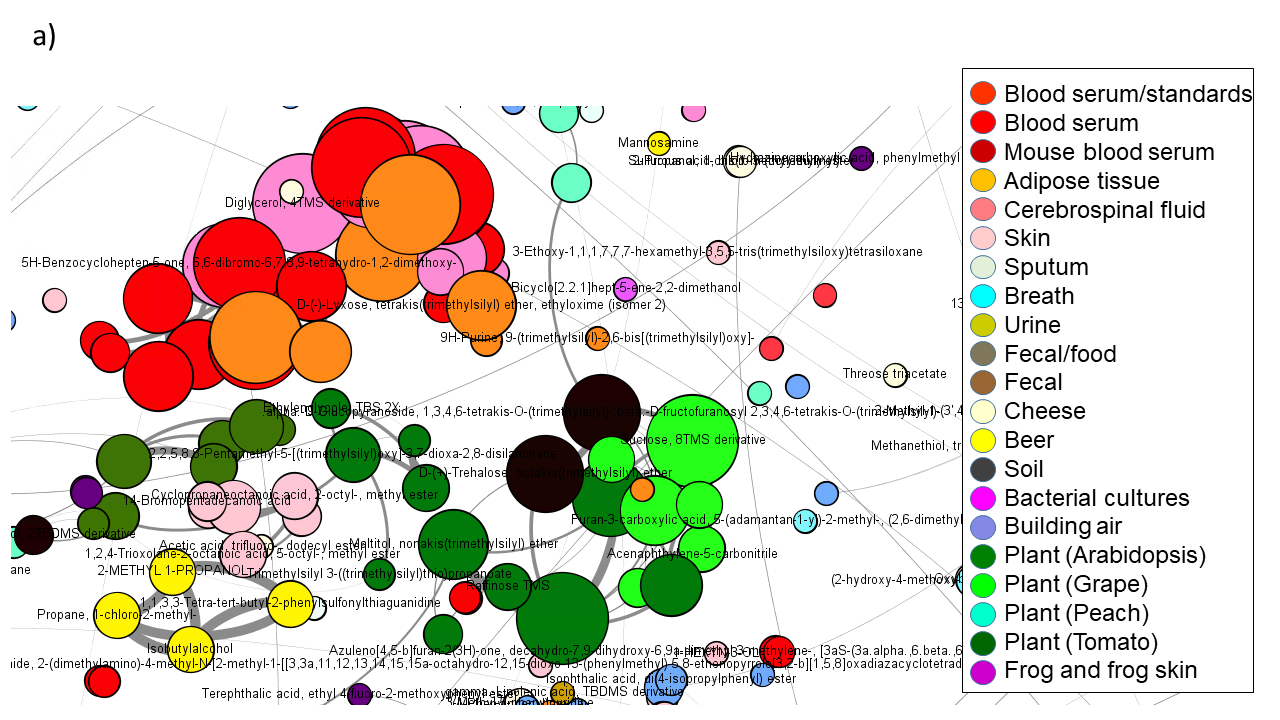
